# Cardiac muscle-restricted partial loss of *Nos1ap* expression has limited impact on electro- and echo-cardiographic features

**DOI:** 10.1101/2022.09.27.509782

**Authors:** Alexa Smith, Dallas Auer, Morgan Johnson, Ernesto Sanchez, Holly Ross, Christopher Ward, Aravinda Chakravarti, Ashish Kapoor

**Affiliations:** Institute of Molecular Medicine, McGovern Medical School, University of Texas Health Science Center at Houston, Houston, Texas 77030; McKusick-Nathans Institute of Genetic Medicine, Johns Hopkins University School of Medicine, Baltimore, Maryland 21205; Department of Molecular Physiology and Biophysics, Baylor College of Medicine, Houston, Texas 77030; Center for Human Genetics and Genomics, New York University School of Medicine, New York, New York 10016.

**Keywords:** Nos1ap, knockout mouse, QT interval, electrocardiogram

## Abstract

Genome-wide association studies (GWAS) of QT interval variation have identified common noncoding variants at the *NOS1AP* gene as the most common genetic regulators of trait variation in the general population. Invoking a *cis*-regulatory mechanistic hypothesis, we have reported identification of a functional enhancer variant underlying the GWAS signal that influenced human cardiac *NOS1AP* expression. Functional studies based on *in vitro* overexpression in murine cardiomyocytes and *ex vivo* knockdown in zebrafish embryonic hearts, by us and others, have demonstrated that NOS1AP expression levels can alter cellular electrophysiology. Here, to explore the role of *NOS1AP* in cardiac electrophysiology at an organismal level, we generated and characterized constitutive and heart muscle-restricted *Nos1ap* knockout mice to assess whether *NOS1AP* disruption alters the QT interval *in vivo*. Constitutive loss of *Nos1ap* led to genetic background-dependent variable lethality at or right before birth. Heart muscle-restricted *Nos1ap* knockouts generated using cardiac specific alpha-myosin heavy chain promoter-driven tamoxifen-inducible Cre resulted in tissue-level *Nos1ap* expression reduced by half. This partial loss of expression had no detectable effect on the QT interval, but led to a small yet significant reduction in the QRS interval. Given that challenges associated with defining the end of T wave on murine electrocardiogram can limit identification of subtle effects on QT interval, and that common noncoding *NOS1AP* variants are also associated with QRS interval, our findings support the role of *NOS1AP* in regulation of the cardiac electrical cycle.

## Introduction

The electrocardiographic QT interval, an index of ventricular repolarization (1), is a clinically relevant, heritable quantitative trait associated with cardiovascular disease mortality in the general population (2). Prolongation or shortening of the QT interval, owing to underlying pathology, genetic variants, or adverse drug reactions, can lead to life-threatening arrhythmias and sudden cardiac death (SCD). QT interval, the time between the start of the Q wave and the end of the T wave in an electrocardiogram (ECG), is itself genetically influenced with a heritability of 35% but also correlated with age, sex and heart rate (3). Beyond the rare, high penetrance, coding mutations leading to phenotypic extremes of QT interval in subjects with Mendelian long QT (LQTS), short QT, or Brugada syndromes (4, 5), common genetic variation is a major source of QT interval variation in the general population. Genome-wide association studies (GWAS) of QT interval have mapped at least 35 common variants-based loci (6–9) contributing to its heritability, and, among them, the locus with the largest contribution (~1.5% of phenotype variation) includes the gene *NOS1AP* on chromosome 1q. Although, the functional role of *NOS1AP* in cardiac repolarization is not established, QT interval associated common variants at the *NOS1AP* locus are also associated with increased risk of SCD in the general population (10, 11) and are genetic modifiers of cardiac outcomes in subjects with LQTS (12). Furthermore, QT interval associated *NOS1AP* variants are also associated with QRS interval, representing ventricular depolarization on an ECG, but the same variants often act in the opposite direction (9, 13).

NOS1AP (<u>n</u>itric <u>o</u>xide <u>s</u>ynthase <u>1</u> <u>a</u>daptor <u>p</u>rotein), initially called CAPON (<u>ca</u>rboxy-terminal <u>P</u>DZ ligand <u>o</u>f <u>n</u>euronal NOS), is the C-terminal PDZ ligand of NOS1/nNOS and was originally cloned from a rat hippocampal cDNA library (14). Nearly all biochemical characterization of *NOS1AP* function so far has been in *in vitro* and *ex vivo* systems, largely based on neuronal cell lines and tissue lysates (15), with very limited knowledge about its cardiac function. A possible relationship between the nitric oxide synthase pathway and cardiac repolarization was not recognized before the QT interval GWAS mapping (6). Invoking a *cis*-regulatory mechanistic hypothesis, we identified a functional enhancer variant underlying the QT interval GWAS signal that influenced *NOS1AP* cardiac transcript expression (16). We also demonstrated that overexpression of long and short isoforms of human *NOS1AP* in neonatal rat ventricular myocytes (NRVMs) led to shortened action potential duration (APD), a potential cellular correlate of QT interval (16). Similarly, others showed that overexpression of *Nos1ap* in guinea pig ventricular myocytes led to reduced APD via inhibition of L-type calcium channels and activation of delayed rectifier potassium channels (17). In contrast, optical mapping of excised whole hearts from developing zebrafish embryos with morpholino-based knockdown of *nos1ap* showed shortened APD (18). Although, directionally inconsistent, these studies indicated that *NOS1AP* expression levels can alter cellular electrophysiology. The differences observed could be simply due to differences in model systems or reflect that both gain and loss of *NOS1AP* mis-regulate its functional complexes.

In an effort to explore the role of *NOS1AP* in cardiac electrophysiology at an organismal level, and to assess whether *NOS1AP* disruption alters the QT interval *in vivo*, a critical knowledge gap, here we have generated and characterized constitutive and heart muscle-restricted *Nos1ap* knockout mice. In this paper, we report that constitutive loss of *Nos1ap* leads to near-complete lethality at or right before birth, that constitutive loss of *Nos1ap* has no major impact on the embryonic heart transcriptome, that heart muscle-restricted *Nos1ap* knockouts generated using cardiac specific alpha-myosin heavy chain promoter-driven tamoxifen-inducible Cre (19) reduces tissue-level *Nos1ap* expression by half, and that this partial loss of *Nos1ap* cardiac expression has no detectable effect on the QT interval, but leads to a small and significant reduction in the QRS interval.

## Results

### Generation of constitutive and conditional Nos1ap knockout mice

Starting with mouse embryonic stem (ES) cells targeting exon 4 of *Nos1ap* (NM_001109985) with the ‘knockout-first allele’ (20), purchased from the Knockout Mouse Project (KOMP) repository, reporter-tagged *Nos1ap* knockout mice with conditional potential (*Nos1ap*^+/tm1a^) were generated by blastocyst injection and germline transmission. The flexible design of the tm1a allele was exploited to generate non-conditional reporter-tagged knockout (reporter-tagged and exon 4-deleted; *Nos1ap*^+/tm1b^), Cre-recombinase conditional knockout (exon 4-floxed; *Nos1ap*^+/tm1c^ or *Nos1ap*^+/fl^) and constitutive null (exon 4-deleted; *Nos1ap*^+/tm1d^ or *Nos1ap*^+/−^) mice by crossing with CMV-Cre (21) and ACTB-Flpe (22) mice (Figure 1). Targeted (tm1a) and derived (tm1b, tm1c and tm1d) alleles were maintained in the C57BL/6J background. We used *Nos1ap*^+/tm1c^ and *Nos1ap*^+/tm1d^ mice from backcross generation number 10 (N10) or beyond in phenotypic studies. Alleles tm1a and tm1b are expected to be Nos1ap protein null by design due to the insertion of a transgene cassette containing the Engrailed 2 splice acceptor, internal ribosome entry site, *lacZ* open reading frame and polyadenylation signal between exons 3 and 4. Allele tm1d is also expected to be Nos1ap protein null due to exon 4 deletion-mediated frame-shift that creates a premature termination codon leading to nonsense-mediated mRNA decay. Western blotting of adult brain cortex tissue lysates showed complete absence of Nos1ap protein in *Nos1ap*^tm1a/tm1a^ and *Nos1ap*^tm1b/tm1b^ mice, and a considerable decrease in protein levels in *Nos1ap*^+/tm1a^ and *Nos1ap*^+/tm1b^ mice compared to wildtype (Figure 2). Similarly, compared to wildtype, *Nos1ap* transcript expression in adult brain cortex tissue as measured by real-time quantitative PCR (qPCR) on cDNA was reduced to 73% (*P*=0.01) in *Nos1ap*^+/tm1a^ and *Nos1ap*^+/tm1b^ mice and to 24% (*P*=7.88×10^−8^) in *Nos1ap*^tm1a/tm1a^ and *Nos1ap*^tm1b/tm1b^ mice (Figure 2; Figure S1). The corresponding values for *Nos1ap* transcript expression in adult left ventricle tissue were 79% (*P*=0.01) and 24% (*P*=8.47×10^−6^) (Figure 2; Figure S1).

**Figure 1:**
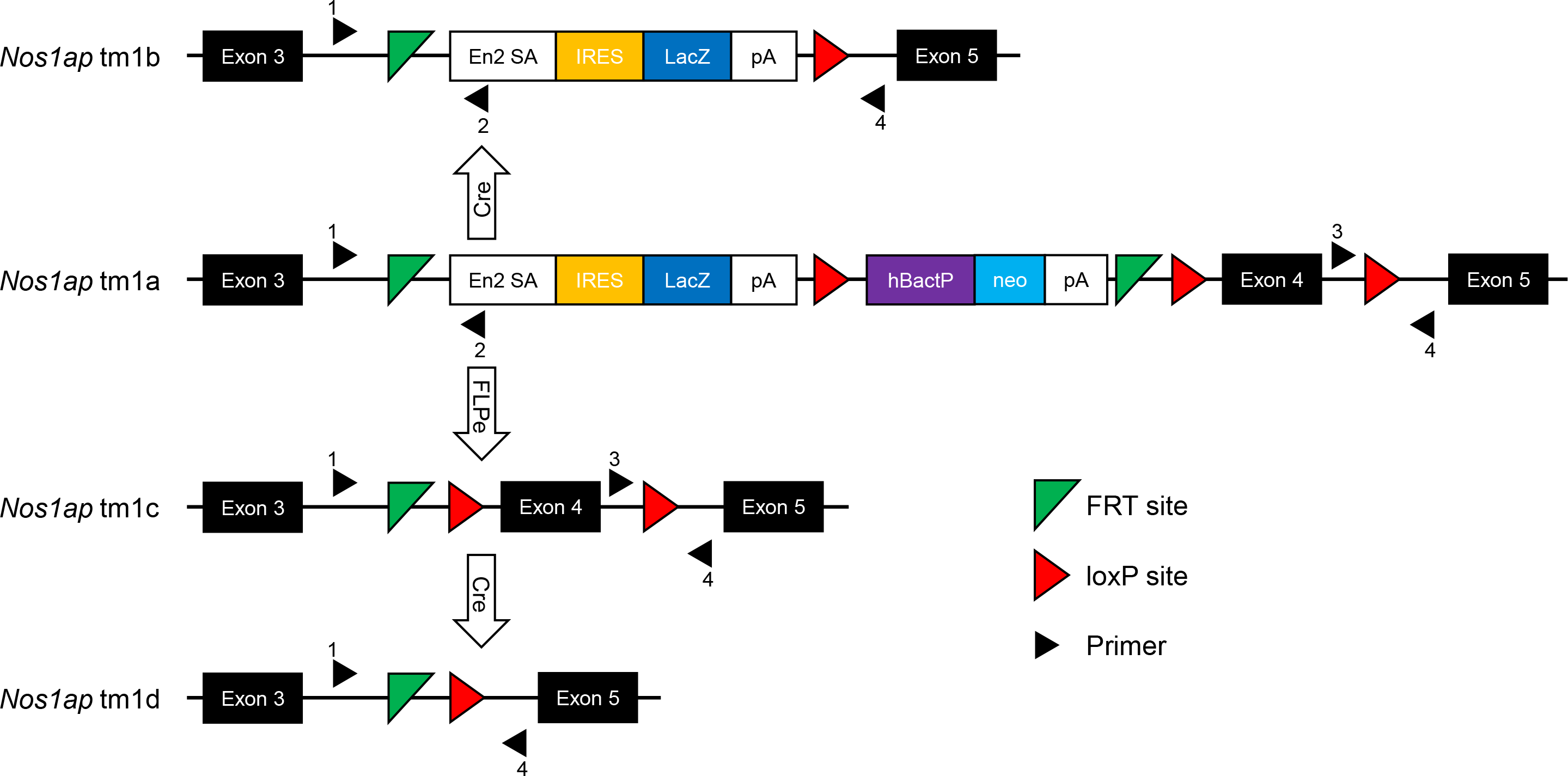
Targeted and derived *Nos1ap* alleles. The ‘knockout-first’ (tm1a) allele contains an internal ribosome entry site (IRES):*lacZ* trapping cassette and a floxed human beta actin (hBactP) promoter-driven *neo* cassette inserted into intron 3 of *Nos1ap*. A splice acceptor sequence from Engrailed 2 (En2 SA) and poly-A (pA) transcription termination signals disrupt *Nos1ap* expression while expressing *lacZ* under the control of the endogenous promoter. Exposure to Cre recombinase mediates conversion of tm1a to tm1b allele to generate a non-conditional *lacZ*-tagged null allele without the *neo* cassette and without the critical region (Exon 4). Exposure to FLPe recombinase mediates conversion of tm1a to tm1c allele to generate a conditional floxed allele, which on further exposure to Cre recombinase can generate either the constitutive null allele (tm1d) or a tissue-restricted *Nos1ap* knockout. Black-filled triangles, numbered 1 to 4, indicate primers used for PCR genotyping; see Tables S1 and S2 for details.

**Figure 2:**
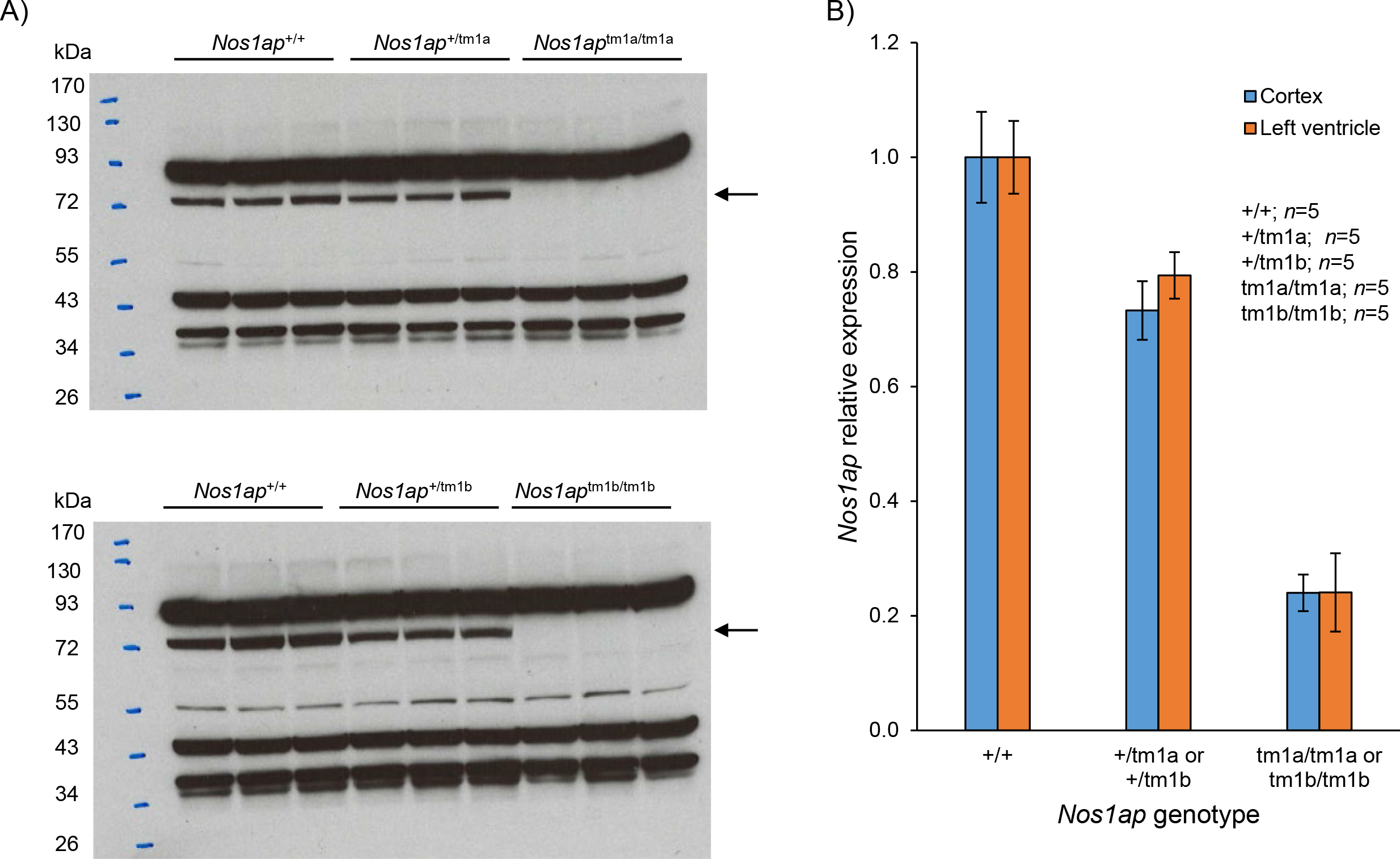
Loss of Nos1ap protein and transcript in *Nos1ap* tm1a and tm1b allele carriers. *A*) Western blot of brain cortex lysates from adult mouse using rabbit polyclonal NOS1AP antibody shows complete absence of Nos1ap protein (immunoreactive band indicated by arrow) in tm1a (*top*) and tm1b (*bottom*) homozygotes, and reduced protein levels in tm1a (*top*) and tm1b (*bottom*) heterozygotes as compared to wildtype mice. Other immunoreactive bands likely indicating non-specific binding were also observed. *B*) Compared to wildtype mice, tm1a or tm1b heterozygotes and tm1a or tm1b homozygotes had significantly reduced *Nos1ap* transcript expression in adult brain cortex and left ventricle tissues. Error bars: SEM.

### Constitutive loss of Nos1ap leads to near-complete lethality

To generate homozygous *Nos1ap* constitutive null mice, free of *lacZ* and *neo* cassette, *Nos1ap*^+/tm1d^ mice were intercrossed. With increasing backcross generation numbers, *Nos1ap*^+/tm1d^ intercrosses were performed at two different locations; first at Johns Hopkins University (JHU) and then at University of Texas Health Science Center at Houston (UTHealth). Among *Nos1ap*^+/tm1d^ intercrosses at JHU (parental mice from N8 to N11), 9.6%, 7.5% and 25.2% of all mice were *Nos1ap*^tm1d/tm1d^ at weaning (P21), birth (P0) and embryonic day 13.5 (E13.5), respectively (Table 1). Among *Nos1ap*^+/tm1d^ intercrosses at UTHealth (parental mice from N14 to N15), 3.4% of all mice were *Nos1ap*^tm1d/tm1d^ at P21 (Table 1). With an expectation of 25% mice being null homozygotes under Mendelian segregation, a significant drop in counts observed at P21 (*P*=7.61×10^−7^ JHU; *P*=1.53×10^−5^ UTHealth), P0 (*P*=1.04×10^−4^), but not at E13.5 (*P*=0.09) (Table 1), indicate that complete loss of *Nos1ap* in *Nos1ap*^tm1d/tm1d^ mice leads to near-complete lethality at birth or during late gestation (after E13.5).

**Table 1:** Genotype distribution from *Nos1ap* constitutive null intercrosses.

|  | **E13.5 (%)<sup>1</sup>** | **P0 (%)<sup>1</sup>** | **P21 (%)<sup>1</sup>** | **P21 (%)<sup>2</sup>** |
| --- | --- | --- | --- | --- |
| +/+ | 37 (33.3) | 35 (32.7) | 70 (32.0) | 31 (35.6) |
| +/- | 46 (41.4) | 64 (59.8) | 128 (58.4) | 53 (60.9) |
| -/- | 28 (25.2) | 8 (7.5) | 21 (9.6) | 3 (3.4) |
| $\chi^2$ | 4.71 | 17.75 | 28.18 | 22.17 |
| *P* | 0.09 | 1.04×10<sup>-4</sup> | 7.61×10<sup>-7</sup> | 1.53×10<sup>-5</sup> |

<sup>1</sup>at JHU; <sup>2</sup>at UTHealth

### Constitutive loss of Nos1ap has no significant impact on E13.5 heart transcriptome

To evaluate the *molecular* consequences of *Nos1ap* loss that might lead to embryonic lethality, we assessed differential gene expression between E13.5 heart transcriptomes from wildtype and *Nos1ap*^tm1d/tm1d^ mice. We performed stranded mRNA-seq (23) in E13.5 heart tissues dissected from five wildtype and five *Nos1ap*^tm1d/tm1d^ male mice. Paired-end 100 bp sequencing was performed on an Illumina HiSeq 2500 with a desired depth of ~30 million paired-end reads per sample. Low quality bases and Illumina adapter sequences were removed from the generated reads using Trimmomatic (24). RSEM (25) was used for estimation of gene and isoform expression level that included mapping of trimmed paired-end reads to the mouse genome and GENCODE transcripts using STAR aligner (26). Gene-level read count data from five wildtype and five mutants were compared to assess differential gene expression using DESeq (27). At a false discovery rate (FDR) of 1% and an absolute log_2_-fold change over 1, only two genes (*Mt2* and *Gm7694*) were differentially expressed (DatasetS1; Figure S2), beyond that expected at *Nos1ap*. Compared to wildtype mice, *Nos1ap* expression was significantly reduced (0.3×; *P*=2.9×10^−31^), *Mt2* expression was significantly increased (2.3×; *P*=4.2×10^−8^) and *Gm7694* expression was significantly reduced (0.4×; *P*=8.5×10^−39^) in *Nos1ap*^tm1d/tm1d^ mice. These findings indicate that constitutive loss of *Nos1ap* has no major widespread impact on the E13.5 heart transcriptome.

### Cardiac muscle-restricted, αMHC promoter-driven tamoxifen-inducible Cre leads to partial loss of expression in Nos1ap floxed mice

Given the near-complete lethality observed in *Nos1ap* constitutive null homozygous mice at birth or during late gestation, we evaluated the effects of cardiac muscle-restricted *Nos1ap* loss using the *Nos1ap* floxed mice and cardiac alpha-myosin heavy chain (αMHC) promoter-driven tamoxifen-inducible Cre recombinase (MerCreMer) transgenic mice (αMHC-MerCreMer) (19). Crosses were setup between *Nos1ap*^+/fl^; +/+ and *Nos1ap*^+/fl^; +/*Tg*^αMHC-MerCreMer^ mice to generate *Nos1ap*^+/+^, *Nos1ap*^+/fl^ and *Nos1ap*^fl/fl^ mice with and without αMHC-MerCreMer transgene. All six genotypes were observed at expected proportions at weaning (*P*=0.75; Table S3). Tamoxifen intraperitoneal (IP) injections were performed in four weeks old animals for five consecutive days at a dosage of 20mg/kg per day to induce Cre-recombinase activity (19), and was followed by a one week waiting period before any experimentation. *Nos1ap*^+/+^, *Nos1ap*^+/fl^ and *Nos1ap*^fl/fl^ mice with and without αMHC-MerCreMer were viable, normal in size, and did not display any gross physical or behavioral abnormalities (data not shown). PCR genotyping of genomic DNA isolated from left ventricle and tail (control) tissue in a subset of animals showed cardiac-restricted excision of the floxed allele, with no ‘leaky’ Cre-recombinase activity in tail tissue (Figure 3; Figure S3). As *αMHC* promoter activity is mostly limited to cardiomyocytes, the unexcised floxed allele PCR band derived from other cell types in left ventricle tissue was also observed in *Nos1ap*^fl^ and *Tg*^αMHC-MerCreMer^ carriers (Figure 3). The effect of Cre-mediated, floxed allele excision on *Nos1ap* transcript expression was evaluated by real-time qPCR using cDNA generated from left ventricle tissue harvested at terminal euthanasia. Among αMHC-MerCreMer transgene positive animals, compared to wildtype (*n*=6), *Nos1ap* expression in left ventricle tissue was reduced to 80% (*P*=0.09) and 52% (*P*=1.85×10^−4^) in *Nos1ap*^+/fl^ (*n*=6) and *Nos1ap*^fl/fl^ (*n*=6) mice, respectively (Figure 3). Among αMHC-MerCreMer transgene negative animals, no significant difference in *Nos1ap* left ventricle expression was observed among *Nos1ap*^+/+^, *Nos1ap*^+/fl^ (*P*=0.91) and *Nos1ap*^fl/fl^ (*P*=0.50) mice (Figure S4).

**Figure 3:**
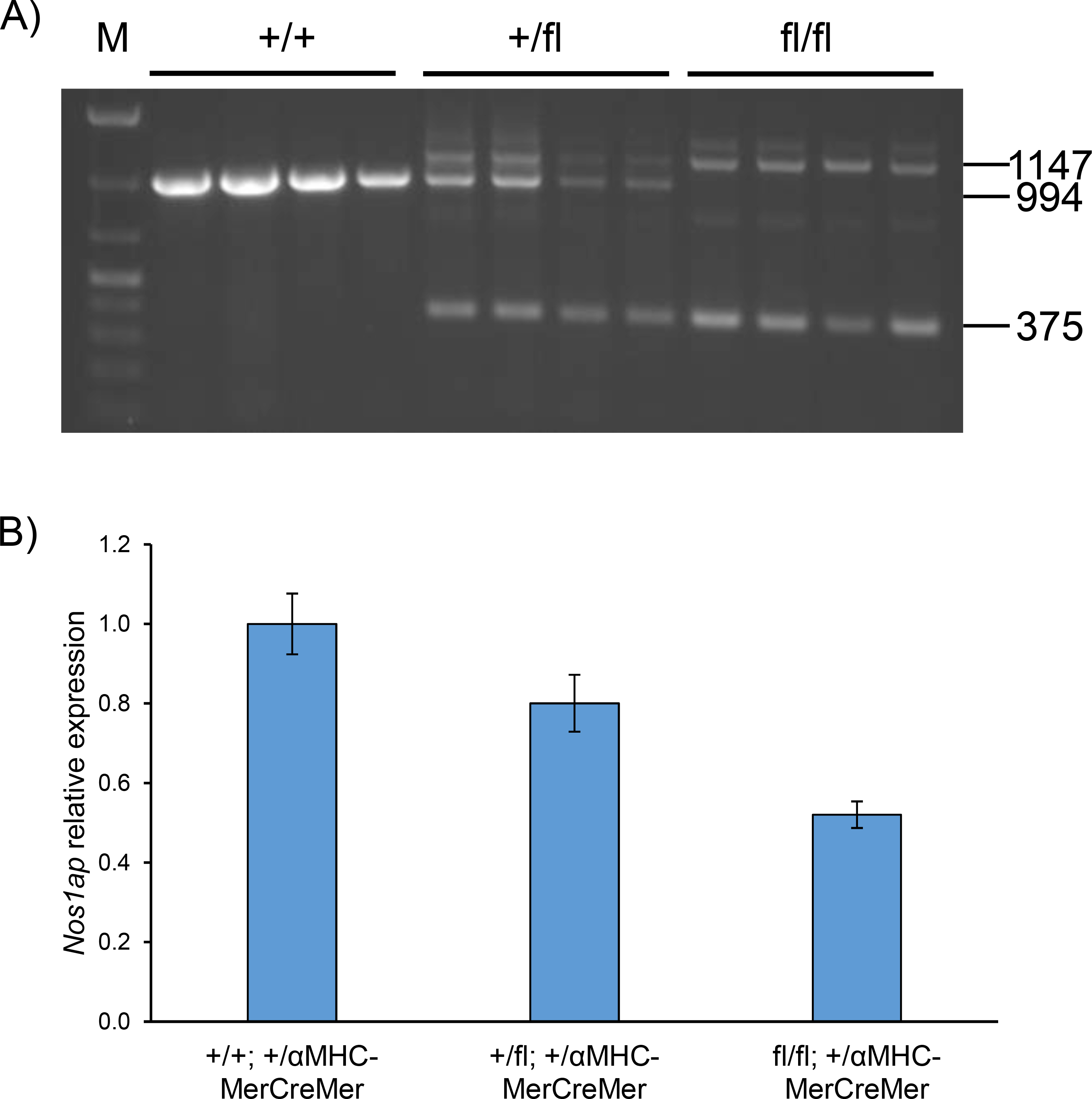
Cardiac αMHC-promoter driven tamoxifen-inducible Cre recombinase leads to excision of floxed allele and loss of *Nos1ap* expression in left ventricle tissue. *A*) Following tamoxifen intraperitoneal injections to induce Cre recombinase activity, agarose gel electrophoresis of amplicons generated by PCR at the *Nos1ap* locus using left ventricle tissue genomic DNA from *Nos1ap*^+/+^, *Nos1ap*^+/fl^ and *Nos1ap*^fl/fl^ mice, all with tamoxifen-inducible αMHC-MerCreMer transgene, shows wildtype (+) allele amplicon (994 bp), floxed (fl) allele amplicon (1147 bp), and floxed excision allele amplicon (375 bp). M: DNA ladder. *B*) Compared to wildtype mice, excision of the *Nos1ap* floxed allele leads to reduced *Nos1ap* transcript expression in left ventricle tissue of floxed heterozygotes and homozygotes.

### Partial loss of Nos1ap cardiac expression impacts QRS interval

Starting at six weeks of age, a week after tamoxifen IP injections, paw contacts-based awake electrocardiogram (ECG) recordings were carried out every two weeks until 24 weeks in *Nos1ap*^+/+^, *Nos1ap*^+/fl^ and *Nos1ap*^fl/fl^ mice, with and without αMHC-MerCreMer transgene, and every four weeks thereafter until 48 weeks in *Nos1ap*^+/+^, *Nos1ap*^+/fl^ and *Nos1ap*^fl/fl^ mice with αMHC-MerCreMer transgene. At each time point, ECGs were recorded in 10 or more animals (in nearly equal sex ratios) for each of the six genotypes (Table S4), and age- and genotype-dependent effects on heart rate-corrected QT intervals (QTc) (28) and other ECG parameters were assessed using linear regression. Overall, based on the predictor variables used (age in weeks, *Nos1ap*^+/fl^ and *Nos1ap*^fl/fl^), the variance explained for QTc remained low (adjusted *R*^2^ of 4.35% and 2.95% with and without αMHC-MerCreMer set; Tables S5, S6). Besides small, but significant age-dependent effects (*β*=0.03, *P*=2.48×10^−19^ in with the αMHC-MerCreMer set, Table S5; *β*=0.04, *P*=2.11×10^−5^ in without the αMHC-MerCreMer set, Table S6), relative to wildtype, absolute genotype-dependent effects from *Nos1ap*^+/fl^; +/ αMHC-MerCreMer (*β*=0.1 ms) or *Nos1ap*^fl/fl^; +/ αMHC-MerCreMer (*β*=0.31 ms) genotypes (Table S5) and *Nos1ap*^+/fl^; +/+ (*β*=0.65 ms) or *Nos1ap*^fl/fl^; +/+ (*β*=0.60 ms) genotypes (Table S6) were negligible. These data indicate that loss of *Nos1ap* cardiac expression had no major impact on ventricular repolarization. None of the other ECG parameters assessed (RR, PR and QRS intervals) varied significantly across genotypes (data not shown).

Given that the paw contacts-based awake ECG recordings (above) may fail to detect subtle effects due to reduced measurement sensitivity, and to collect longer ECG recordings to evaluate the effect of an acute pharmacological challenge, we transitioned to ECG measurements in anesthetized animals using surface electrodes. The set of animals with 48 weeks age awake ECG recording above (*Nos1ap*^+/+^, *Nos1ap*^+/fl^ and *Nos1ap*^fl/fl^ with αMHC-MerCreMer; Table S4) were evaluated further by ECG and echocardiography under anesthesia, followed by terminal euthanasia to harvest tissues for gene expression studies. Surface ECG recordings were performed at baseline and after IP injections of isoproterenol, a beta-adrenergic agonist, at 1mg/kg and 5mg/kg doses. However, no significant differences in QTc were observed from surface ECG recordings at baseline and under stress across the three genotypes (Figure 4), again indicating lack of a genotype-dependent effect on ventricular repolarization. None of the other ECG parameters differed significantly across the three genotypes (Figure S5), except a small, but consistent trend of reduced QRS interval in *Nos1ap* floxed homozygotes (mean QRS interval 8.49ms vs. 7.85ms at baseline (*P*=0.05) and 10.09ms vs. 9.21ms under stress (*P*=0.08) for 1mg isoproterenol in wildtype and floxed homozygotes, respectively; 8.32ms vs. 7.79ms at baseline (*P*=0.06) and 10.01ms vs. 8.95ms under stress (*P*=0.04) for 5mg isoproterenol in wildtype and floxed homozygotes, respectively; 8.40ms vs. 7.82ms at baseline (*P*=0.005) and 10.05ms vs. 9.07ms under stress (*P*=0.005) after combining 1mg and 5mg isoproterenol data in wildtype and floxed homozygotes, respectively; Figure 4). A linear regression model using condition (baseline or under stress), isoproterenol dose (1mg or 5mg), and genotype (wildtype or floxed homozygote) as predictors, explained 45.06% of the observed variance in QRS intervals (*P*=2.51×10^−11^), with genotype having a significant effect (*β*=−0.78, *P*=1.25×10^−4^; Table S7).

**Figure 4:**
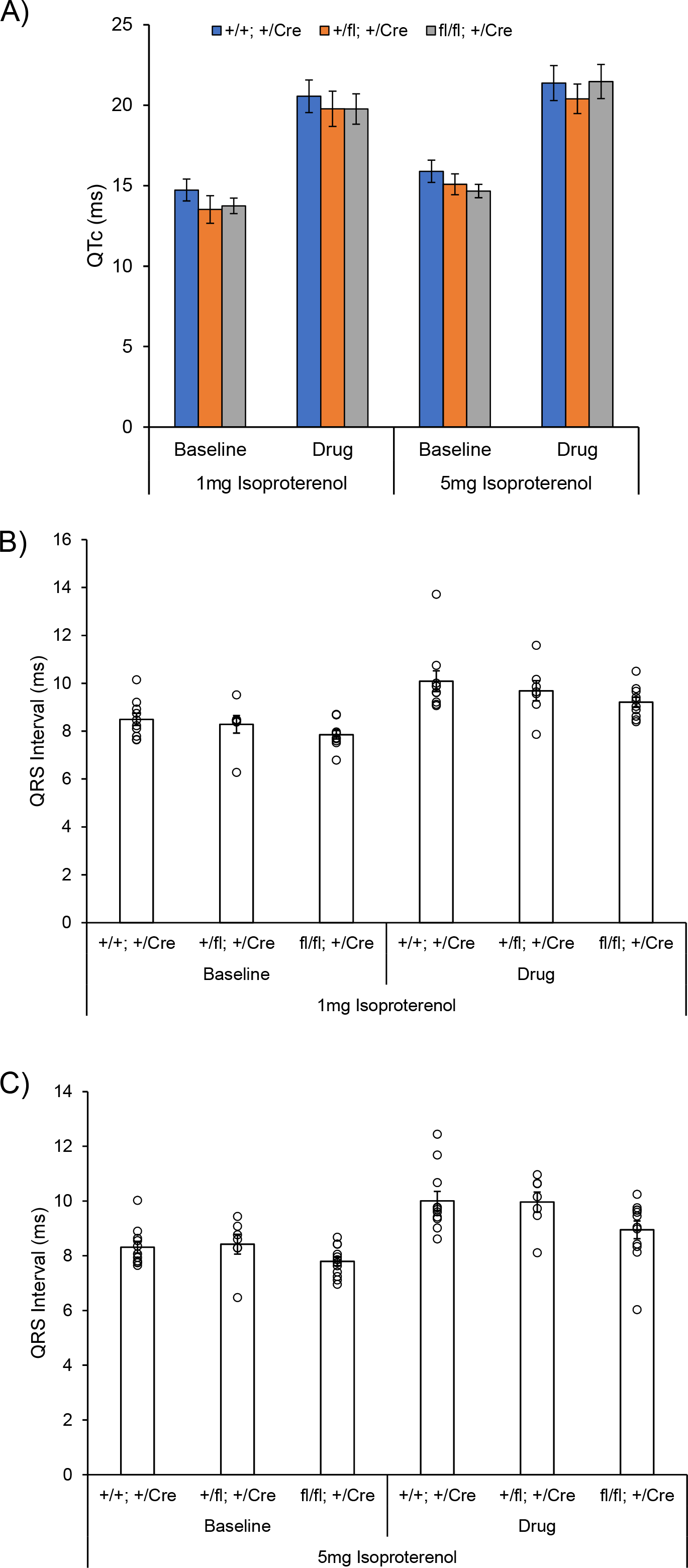
Cardiac muscle-specific loss of *Nos1ap* expression reduces QRS interval without a significant impact on QT interval. *A*) Mean heart rate-corrected QT interval (QTc) from ECG recording under anesthesia at baseline and after injecting 1mg/kg or 5mg/kg body weight doses of isoproterenol in *Nos1ap*^+/+^, *Nos1ap*^+/fl^ and *Nos1ap*^fl/fl^ mice, all with tamoxifen-inducible αMHC-MerCreMer transgene, shows no significant difference across genotypes. Error bars: SEM. *B*) Mean (*bar chart*) and *C*) individual (*dot plot*) QRS intervals from ECG recording under anesthesia at baseline and after injecting 1mg/kg (*B*) or 5mg/kg (*C*) body weight doses of isoproterenol in *Nos1ap*^+/+^, *Nos1ap*^+/fl^ and *Nos1ap*^fl/fl^ mice, all with tamoxifen-inducible αMHC-MerCreMer transgene, shows a small, but significant reduction in QRS interval in floxed homozygotes. Error bars: SEM.

To evaluate potential effects on heart structure and function, echocardiography was performed in anesthetized animals, where short axis image of left ventricles were acquired in M-mode and analyzed for structure and function outcome measures. Besides a trend of reduced ejection fraction (EF) and fractional shortening (FS) observed only in *Nos1ap* floxed allele carrier females (wildtype vs floxed heterozygote: *P*=0.01 for EF and *P*=0.01 for FS; wildtype vs. floxed homozygote: *P*=0.03 for EF and *P*=0.03 for FS) (Figure S6), none of the other echocardiographic parameters differed significantly across the three genotypes (data not shown), indicating that loss of *Nos1ap* cardiac expression had no major impact on left ventricle structure and function.

## Discussion

Following the GWAS mapping of QT interval inter-individual variation near *NOS1AP* (6-9), and subsequent *in vitro* and *ex vivo* expression perturbation studies showing that *NOS1AP* expression levels influence APD (16–18), our goal here, using gene knockout mouse models, was to assess if loss of *Nos1ap* expression *in vivo* impacts QT interval and other electrocardiographic features. Although, no significant impact on QT interval was identified in heart muscle-restricted *Nos1ap* knockout mice, a small yet significant reduction in QRS interval was observed, supporting the role of *NOS1AP* in regulation of the cardiac electrical cycle. It is important to note that although the effect of common *NOS1AP* variants is largest for QT interval, the same variants also show an attenuated association with QRS interval (9, 13), surprisingly in the opposite direction. The QRS interval, representing ventricular depolarization, is also known to be positively correlated with the QTc interval (*r*=0.44), which represents ventricular repolarization (29). Overall, across all ECG measurements in conscious and anesthetized mice here, QRS and QTc intervals were positively correlated (*r*=0.30 and *r*=0.74 in awake and anesthetized mice, respectively). Furthermore, due to differences in ionic currents that generate different shapes of ventricular action potential in human and mouse (1), the ST segment is lacking in mice and the amplitude of T wave on mouse ECG is relatively small, to the extent that existence of an actual T wave on mouse ECG is a matter of longstanding debate (30). The small amplitude of T wave makes pinpointing its end, determined as the return of signal to the voltage corresponding to the mean isoelectric value, error-prone, and this reduced measurement accuracy can lead to failures in detecting subtle effects (31). Thus, even though *Nos1ap* expression levels have been shown to alter APD in rat and guinea pig cardiomyocytes *in vitro* (16, 17), detecting that effect at an organismal level as altered QT interval on mouse ECG is challenging (31). In addition, due to the lack of a plateau in mouse cardiac action potentials, the QRS complex on mouse ECG corresponds to the spread of ventricular depolarization and the early phase of ventricular repolarization (31). Therefore, it is possible that the reduced QRS interval we are observing in heart muscle-restricted *Nos1ap* knockout mice is indicative of shortened ventricular repolarization, at least in the early phase. This observation aligns (in terms of directionality) with the optical mapping-based reduced APD reported from zebrafish excised hearts with morpholino-based *nos1ap* knockdown (18). As outlined below, there are other plausible reasons for not finding an overt QT interval phenotype in our *Nos1ap* knockout mice, in addition to asking if a gene knockout is always an appropriate model to characterize its function.

Constitutive loss of *Nos1ap* in null homozygous mice (*Nos1ap*^tm1d/tm1d)^ led to near-complete lethality at a time-point after E13.5, indicating an essential role in late embryonic development, organ maturation or the process of birth. It remains unknown why this lethal phenotype was not fully-penetrant, but there was a trend of increased penetrance as the backcross generation number for the heterozygous mice used in the intercrosses increased from N8 to N15 (9.6% vs. 3.4% null homozygotes at weaning). Constitutive loss of *Nos1ap* in reporter-tagged knockout with conditional potential (*Nos1ap*^tm1a/tm1a^) and in non-conditional reporter-tagged knockout (*Nos1ap*^tm1b/tm1b^) also displayed near-complete lethality (7.7% and 8.8% null homozygotes at weaning for tm1a and tm1b, respectively) from intercrosses at earlier backcross generation numbers (N2 to N4 for tm1a and N5 to N6 for tm1b). Taken together, across all three null alleles and backcross generation numbers (N2 to N15), 8.0% of mice at weaning from intercrosses were null homozygotes, underscoring the embryonic or early postnatal lethality led by constitutive loss of *Nos1ap*. These observations are in sharp contrast with the publicly available viability data from the International Mouse Phenotyping Consortium (IMPC; https://www.mousephenotype.org/) for *Nos1ap*^+/tm1b^ intercrosses that show expected Mendelian ratios for the three genotypes. A pure genetic background of our mice (C57BL/6J) versus the mixed genetic backgrounds of IMPC mice at the levels of test-cross (germline transmission), tm1a to tm1b conversion (Cre-driver), and maintenance of alleles could potentially explain this contrast, emphasizing the effects genetic backgrounds can have on gene function and phenotypes.

Although previous studies have not evaluated a role for *NOS1AP* in gene expression regulation, we explored E13.5 heart transcriptome with the aim to uncover molecular events that may underlie late embryonic or preweaning lethality observed in *Nos1ap* constitutive null homozygotes. However, we did not find any major impact on E13.5 heart transcriptome as assessed by RNA-seq and differential gene expression analysis. Given these data along with the presence of a functional mouse fetal heart before E13.5, we conclude that the cause of lethality is non-cardiac. Indeed, *NOS1AP* is widely expressed in several human and mouse embryonic and adult tissues (16, 32), with highest expression levels in brain tissues. Therefore, assessing the transcriptional molecular consequences of *Nos1ap* loss-of-function in additional tissues/organs remains critical.

We would like to highlight that our study is not the first report of *Nos1ap* knockout mouse model. A previous study reported generation of *Nos1ap* knockout mice and characterization by ECG and echocardiography (33). However, there were no data showing loss of *Nos1ap* transcript and/or protein in mutant mice, which we provide both for the constitutive and tissue-restricted null. The targeted mouse ES cells used in that study carried deletion of *Nos1ap* exon 3, which, based on sequence alone, is expected to generate an in-frame 30 amino acid deletion (full length wildtype protein 503 amino acids) that likely has limited impact on protein structure and function. In contrast, the KOMP generated mouse ES cells (20) we used targeted *Nos1ap* exon 4, deletion of which creates a frame-shift, introducing a premature termination codon leading to nonsense-mediated mRNA decay and complete loss of protein. Moreover, the exon targeted in KOMP ES cells is a ‘critical’ exon common to all transcript variants that, when deleted, creates a frame-shift mutation (20). Also, in contrast to our findings, the *Nos1ap* mutant homozygotes in the C57BL/6J background were reported to be viable. At baseline, no difference between wildtype and *Nos1ap* knockout mice in surface ECG and echocardiography was reported. Lastly, following injection of doxorubicin, a drug known to induce cardiotoxicity (34), the authors did report longer QTc intervals in *Nos1ap* knockout mice. Keeping in mind that no data showing loss of *Nos1ap* expression was reported, at least to us, that conclusion remains debatable as similar changes in QTc intervals and several other physiological measurements were observed in doxorubicin-treated wildtype controls, indicating presence of mostly genotype-independent, drug-induced effects (33).

Since Cre expression in αMHC-MerCreMer transgene is driven by a cardiomyocyte-specific promoter (19), the floxed allele in cardiac endothelial cells, fibroblasts and other stromal cells in *Nos1ap*^fl/fl^; αMHC-MerCreMer mice will remain unexcised. Given the widespread expression of *NOS1AP* in various human and mouse tissues (16, 32), it is likely that *Nos1ap* is also expressed in non-cardiomyocyte cells in heart tissue. It is also possible that Cre-mediated excision of the floxed allele (two copies per cell in *Nos1ap*^fl/fl^) is not complete in all cardiomyocyte nuclei. Alone or together, these scenarios can explain why the loss of *Nos1ap* transcript expression in left ventricle tissue in *Nos1ap*^fl/fl^ was half of that in wildtype mice. In order to assess *Nos1ap* expression at the cell type level in mouse left ventricles, we checked for *Nos1ap* expression in a published single cell RNA-seq (scRNA-seq) dataset from adult (P56) left ventricle tissue (35). However, across the ~2,500 cells sequenced, including both cardiomyocytes- and non-cardiomyocytes-enriched cells, *Nos1ap* expression was not detected in any of the identified cell clusters, most likely due to limited sensitivity (median of 2610 genes/cell) (35), in contrast to its detection by TaqMan Gene Expression assay in bulk tissue.

Although previous *in vitro* studies have implicated cardiomyocyte-based *Nos1ap* effect on cellular electrophysiology, it remains possible that other cell types are involved in the regulation of cardiac electrical cycle. For example, a neuronal effect is possible given that autonomic nervous system plays an important role in modulation of cardiac electrophysiology and arrhythmogenesis (36, 37), and that *Nos1ap* has the highest expression level in nervous system tissues (14, 16). Even when restricting to cardiomyocyte-mediated effects, incomplete deletion of *Nos1ap* floxed allele copies in cardiomyocytes of *Nos1ap*^fl/fl^; αMHC-MerCreMer mice may fail to produce sufficient loss-of-function necessary to induce an overt ECG phenotype. The converse is also possible, where a complete or near-complete *Nos1ap* loss-of-function (knockout) in cardiomyocytes is rescued by yet unknown compensatory pathways (38, 39). The latter raises an important question for the utility of gene knockouts to understand gene function. This is especially applicable for genes uncovered by GWAS of common diseases and traits, as is the case here, where an ideal *in vivo* model to assess variable gene-expression based outcomes should involve an allelic series with target gene expression varying from low to high, as opposed to its complete absence, although generating such an allelic series remains challenging, at least in higher vertebrate model organisms.

## Materials and Methods

### Generation of constitutive and conditional Nos1ap null mice

Targeted mouse ES cells with *Nos1ap* ‘knockout-first allele’ (*Nos1ap*^tm1a^; reporter-tagged insertion with conditional potential) (20) derived from the parental ES cell line JM8A3.N1 (*A/a*; *Tyr*^+/+^) of the C57BL/6N strain (40) were purchased from the KOMP repository. Injections into albino blastocysts (C57BL/6 *a/a*; *Tyr*^c/c^) and generation of G0 chimeras with agouti coat color was performed at Texas A&M Institute of Genomic Medicine. Chimeric mice were crossed to C57BL/6J (*a/a*; *Tyr*^+/+^) and G1 mice were genotyped by PCR (Table S1) to assess germline transmission. *Nos1ap*^+/tm1a^ mice were crossed with CMV-Cre (Jax stock #006054) (21) to generate reporter-tagged knockout mice (*lacZ*-tagged, *neo*^R^-deleted, *Nos1ap*-exon4 deleted; *Nos1ap*^+/tm1b^), and crossed with ACTB-Flpe (Jax stock #005703) (22) to generate Cre-recombinase conditional knockout mice (*lacZ*-deleted *neo*^R^-deleted, *Nos1ap*-exon4 floxed; *Nos1ap*^+/tm1c^ or *Nos1ap*^+/fl^). Subsequently, *Nos1ap*^+/tm1c^ mice were crossed with CMV-Cre (Jax stock #006054) (21) to generate constitutive null mice (*lacZ*-deleted *neo*^R^-deleted, *Nos1ap*-exon4 deleted; *Nos1ap*^+/tm1d^ or *Nos1ap*^+/−^). All alleles were maintained by backcrossing to C57BL/6J, and mice with backcross generation ≥N10 were used for phenotyping of *Nos1ap* tm1c and tm1d alleles. All protocols for animal care, use and euthanasia were reviewed and approved by the Institutional Animal Care and Use Committees at JHU, UTHealth, and Baylor College of Medicine (BCM), and were in accordance with the Association for Assessment and Accreditation of Laboratory Animal Care guidelines. All animals were fed a standard rodent chow *ad libitum*. Genomic DNA was isolated from tail-tips of 3 weeks old mice at weaning or from tail-tips and left ventricle tissue of euthanized adult mice or from E13.5 embryos following standard methods. All mice were genotyped by PCR (see Table S1 for primers and Table S2 for amplicons; PCR conditions available on request). Alleles tm1a, tm1b and tm1d are Nos1ap protein null by design.

### RNA isolation and gene expression analyses

Adult mice were euthanized using inhaled isoflurane in a closed chamber, and dissected tissues were snap-frozen in liquid nitrogen prior to storage in −80°C. Total RNA was extracted from ~20mg dry tissue using TRIzol (Invitrogen, MA) following the manufacturer’s instructions. DNase digestion and RNA clean-up were performed using RNeasy Mini kit and RNase-Free DNase set (Qiagen, MD), following the manufacturer’s instructions. cDNA was synthesized by oligo-dT primed reverse transcription performed on 1μg of total RNA using SuperScript III First-Strand Synthesis System (Invitrogen, MA), following the manufacturer’s instructions. Quantitative expression analysis of *Nos1ap* was performed using mouse-specific TaqMan Gene Expression assay (Mm01290688_m1; mapping to exons 5 and 6) (Applied Biosystems, MA). Real-time qPCR was performed on a 7900HT Fast Real-Time PCR System or QuantStudio 5 Real-Time PCR System (Applied Biosystems, MA) and analyzed using Sequence Detection System Software v.2.1 or QuantStudio Design and Analysis Software v.1.2 (Applied Biosystems, MA). Expression was measured in technical triplicates and the averages of the threshold cycle (C_t_) values were used for analysis. *Actb* expression, assessed using mouse *Actb* Endogenous Control TaqMan Gene Expression assay (Applied Biosystems, MA), was used for normalization.

### Western blotting

Nos1ap expression was evaluated in mouse brain cortex lysates using commercially available rabbit polyclonal NOS1AP antibody (R-300, Santa Cruz Biotechnology, TX). Adult mice were euthanized using inhaled isoflurane in a closed chamber, and dissected tissues were snap-frozen in liquid nitrogen prior to storage in −80°C. Whole tissue protein extracts were prepared by cryogenic pulverization of ~20mg of tissue with Cellcrusher (Cellcrusher, OR). Pulverized tissue was suspended in modified RIPA buffer supplemented with protease inhibitor cocktail (Roche, IN). Following sonication, tissue and cell debris were removed by centrifugation, and protein concentration was determined by Bio-Rad DC Protein assay (Bio-Rad, CA). Samples (75μg) were denatured and analyzed by Western blotting following standard methods (41).

### RNA-seq library preparation, sequencing and analyses

RNA-seq was performed in E13.5 heart tissues from five *Nos1ap*^+/+^ and five *Nos1ap*^−/-^ males. Total RNA was isolated from E13.5 heart tissue using RNeasy Mini Kit following the manufacturers’ recommendations (Qiagen, MD) that included the on-column DNase digestion using RNase-Free DNase set (Qiagen, MD). KAPA Stranded mRNA-Seq kit (KAPA Biosystems, MA) was used to generate indexed Illumina platform sequencing libraries. Briefly, polyA RNA was captured from 1μg total RNA using magnetic oligo-dT beads. After elution from the magnetic beads, polyA RNA was fragmented to generate inserts ranging in size from 100-200 bp, followed by random priming and reverse transcription to generate double-stranded cDNA. Next, after performing a 1.8× SPRI cleanup using AMPure XP beads (Agencourt, IN), dAMP was added to 3’-ends of the cDNA fragments followed by ligation with indexed 3’-dTMP Illumina TruSeq adapters. Ligated fragments were subsequently size selected using PEG/NaCl SPRI solution and underwent PCR amplification (12 cycles) to generate the sequencing libraries. After performing a 1× SPRI cleanup using AMPure XP beads (Agencourt, IN), a sample from each library was used to assess library fragment size distribution by electrophoresis using BioAnalyzer High Sensitivity DNA Assay (Agilent Technologies, CA) and to assess library concentration by qPCR using KAPA library quantification kit (KAPA Biosystems, MA). Equimolar amounts of libraries were pooled and sequenced on an Illumina HiSeq 2500 instrument using standard protocols for paired end 100 bp sequencing with a desired sequencing depth of ~30 million paired-end reads per library. Paired-end read fastq files were quality checked using FASTQC (version: 0.11.5) (http://www.bioinformatics.babraham.ac.uk/projects/fastqc/) and then processed using Trimmomatic (version: 0.36) (24), for removing adapters and other Illumina-specific sequences from the reads, and, for performing a sliding-window based trimming of low quality bases from each read (ILLUMINACLIP:TruSeq3-PE-2.fa:2:30:10:1:TRUE LEADING:3 TRAILING:3 SLIDINGWINDOW:4:15 MINLEN:36). For estimating gene and isoform expression levels, we first extracted reference transcript sequences from the mouse genome (GRCm38, primary assembly) based on the GENCODE (http://www.gencodegenes.org/mouse_releases/current.html) primary assembly gene annotation (release M10) and built STAR aligner (26) indices using the RSEM software package (version: 1.2.31) (25). Trimmed paired-end reads from each sample were then aligned to the reference transcript sequences by calling the STAR aligner within RSEM and using alignment parameters from the ENCODE STAR-RSEM long RNA-seq pipeline (--outSAMunmapped Within --outFilterType BySJout -- outSAMattributes NH HI AS NM MD --outFilterMultimapNmax 20 -- outFilterMismatchNmax 999 --outFilterMismatchNoverLmax 0.04 --alignIntronMin 20 --alignIntronMax 1000000 --alignMatesGapMax 1000000 --alignSJoverhangMin 8 --alignSJDBoverhangMin 1 --sjdbScore 1 --quantMode TranscriptomeSAM). Gene and isoform expression levels were then estimated in each sample from these transcriptome alignments using RSEM, keeping in mind the strandedness of the prepared RNA-seq libraries (--forward-prob 0.0). Gene-level read count data generated by RSEM was compared between wildtype and mutant mice to assess differential gene expression using DESeq (version: 1.24.0) (27). Only those genes where the sum of read counts across the 10 samples was >1, were retained for differential gene expression analysis. Although, release M10 of the GENCODE primary assembly gene annotation has 48,526 genes, we limited differential gene expression comparison to only protein coding genes (22,098). To address multiple hypothesis testing, observed *P*-values were adjusted based on the Benjamini-Hochberg FDR procedure (42, 43). All data have been deposited in NCBI’s GEO and are accessible at GEO Series accession number GSE210266.

### Cardiac muscle-restricted Nos1ap loss of expression

To generate tamoxifen-inducible zero, one or two copy loss of *Nos1ap* in cardiac muscle, we utilized the mouse cardiac specific alpha-myosin heavy chain promoter driven tamoxifen-inducible Cre recombinase transgenic line (αMHC-MerCreMer; Jax stock #005657) (19). *Nos1ap*^+/fl^; +/+ mice were crossed with *Nos1ap*^+/fl^; +/*Tg*^αMHC-MerCreMer^ mice to generate *Nos1ap*^+/+^, *Nos1ap*^+/fl^ and *Nos1ap*^fl/fl^ mice with and without αMHC-MerCreMer transgene. To induce Cre recombinase, four weeks old mice were treated with tamoxifen (Sigma, MO) by IP injection once a day for five continuous days at a dose of 20mg/kg per day (19). Tamoxifen stock solution was prepared weekly by dissolving 50mg tamoxifen in 10ml of corn oil (Sigma, MO) and stored at 4°C. Following a one-week gap post-injections, a small number of mice were euthanized to assess Cre recombinase-mediated deletion of floxed allele by PCR genotyping of tail and heart genomic DNA samples.

### Electro- and echo-cardiographic measurements

ECG in conscious mice were performed using ECGenie System (Mouse Specifics, MA), with data acquisition using LabChart (ADIstruments, CO) and analysis using EzCG Analysis software (Mouse Specifics, MA) following the manufacturer’s instructions. Briefly, animals were placed on the recording platform to acclimate for at least 5 minutes before starting data collection. Data was collected for ~10 minutes at a sampling rate of 2000/s and the following filter settings: 3Hz high pass, 100Hz low pass, 60Hz notch and Mains filter. At least three different segments of ECG signals, each with 20 or more heart beats, were exported and analyzed to report various ECG indices. ECG in anesthetized mice were captured using Rodent Surgical Monitor+ (Indus Instruments, TX) and PowerLab 4/35 (ADInstruments, CO), and analyzed using LabChart 8 software (ADInstruments, CO). Briefly, animals were kept anesthetized by inhaled isoflurane delivered in oxygen (induction at 2.5-4%, maintenance at 1.5% isoflurane) via a nose cone, and placed supine on the monitoring platform with paws taped in contact with electrodes for recording ECG waveforms. The monitoring platform was heated and feedback controlled via rectal thermometer to maintain thermal homeostasis (36-38°C) during the ~25-minute recording session. Following a five-minute baseline recording, isoproterenol hydrochloride (USP, MD) at 1mg/kg or 5mg/kg dose was delivered via IP injection for pharmacological challenge. Post-injection measurement was collected for the next 20 minutes. ECG measurements for the two isoproterenol hydrochloride doses were separated by five days. To facilitate signal analysis, a digital 5Hz high pass filter was applied and the in-built ECG analysis suite was used for identifying ECG beats and analysis of ECG parameters following manufacturer’s instructions. The QT interval was corrected for heart rate as described earlier (28). Two days after the second ECG measurement, echocardiography was performed using a Vevo 2100 system (FUJIFILM VisualSonics, WA) with MS550S transducer, following the manufacturer’s instructions. Briefly, animals were anesthetized by inhaled isoflurane and maintained at a body temperature between 36-38°C as described above. Animals and ultrasound transducer probe were positioned to facilitate short-axis imaging of left ventricle at the level of papillary muscles, and B-mode and M-mode images were acquired. Quantification of left ventricle structure and function from M-mode images was performed using the manufacturer’s software that permitted assessment of ventricle wall thickness, inner diameter and derived measures including EF and FS.

### Statistical analyses

Counts data were compared using *χ*^2^ contingency tests. Student’s *t*-test was utilized for comparing mean values between groups. Linear regression was used to evaluate the effect of multiple predictors on ECG parameters.

## Acknowledgements

This work was supported by funds from McGovern Medical School UTHealth (to A.K.) US NIH grant R01 HL086694 (to A.C.), and the Mouse Metabolism and Phenotyping Core at BCM with funding from NIH (UM1 HG006348, R01 DK114356, R01 HL130249). The mouse strain used for this research project was created from ES cell clones EPD0804_4_E03 and EPD0804_4_G02, obtained from the KOMP repository (www.komp.org) and generated by the CSD consortium for the Knockout Mouse Project (KOMP), with funding from NIH (U01 HG004085, U01 HG004080, U42 RR024244).

## Supplementary Information

**Figure S1:**
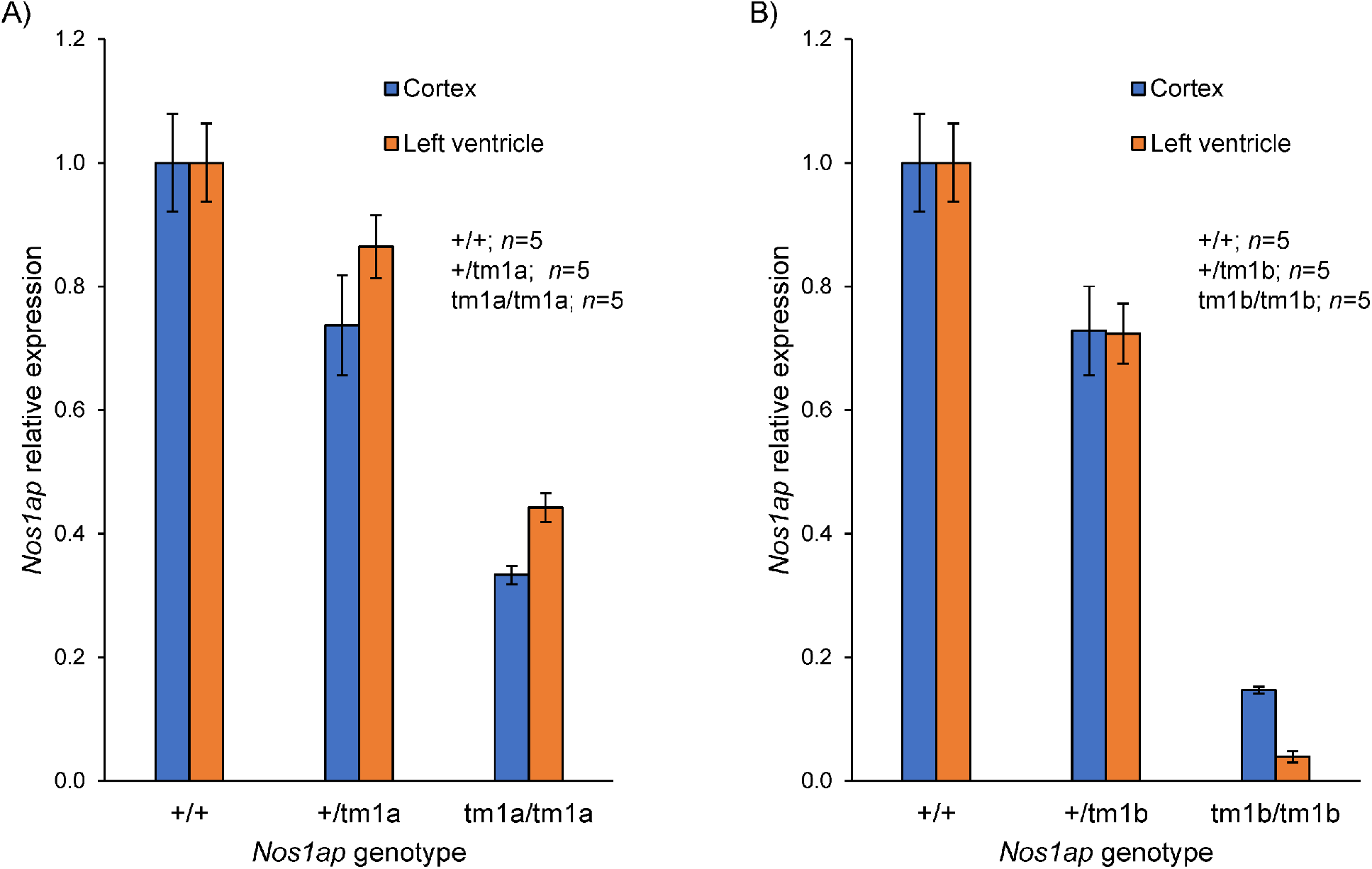
Reduced *Nos1ap* expression in targeted (tm1a) and derived (tm1b) allele carriers. Compared to wildtype mice (+/+), *Nos1ap* tm1a (*A*) and tm1b (*B*) carriers and homozygotes have reduced *Nos1ap* expression in adult brain cortex and left ventricle tissues. *P* values for wildtype to heterozygote comparisons: 0.05 (tm1a cortex), 0.13 (tm1a left ventricle), 0.03 (tm1b cortex), and 0.01 (tm1b left ventricle). *P* values for wildtype to homozygte comparisons: 3.4×10^−5^ (tm1a cortex), 3.5×10^−5^ (tm1a left ventricle), 5.0×10^−6^ (tm1b cortex), 3.9×10^−7^ (tm1b left ventricle). Error bars: SEM.

**Figure S2:**
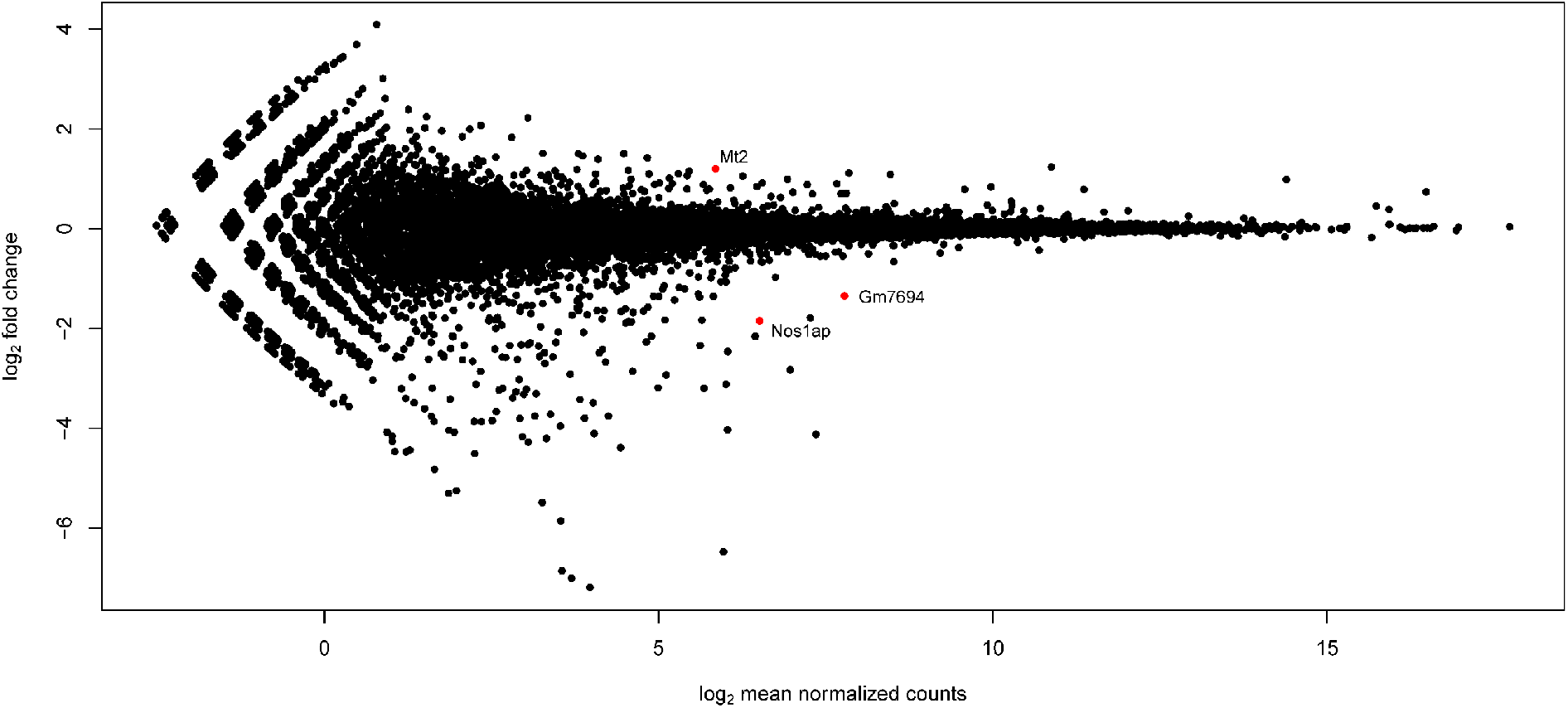
Loss of *Nos1ap* expression has no major effect on E13.5 heart transcriptome. Volcano plot showing log_2_ mean normalized read counts (X-axis) and log_2_ fold change (Y-axis) comparing the E13.5 heart transcriptome of *Nos1ap*^−/-^ to wildtype mice. Each dot represents a gene. Red dots indicate the three genes that are differentially expressed in mutant mice with FDR <1% and absolute log_2_ fold change >1, and black dots indicate other genes.

**Figure S3:**
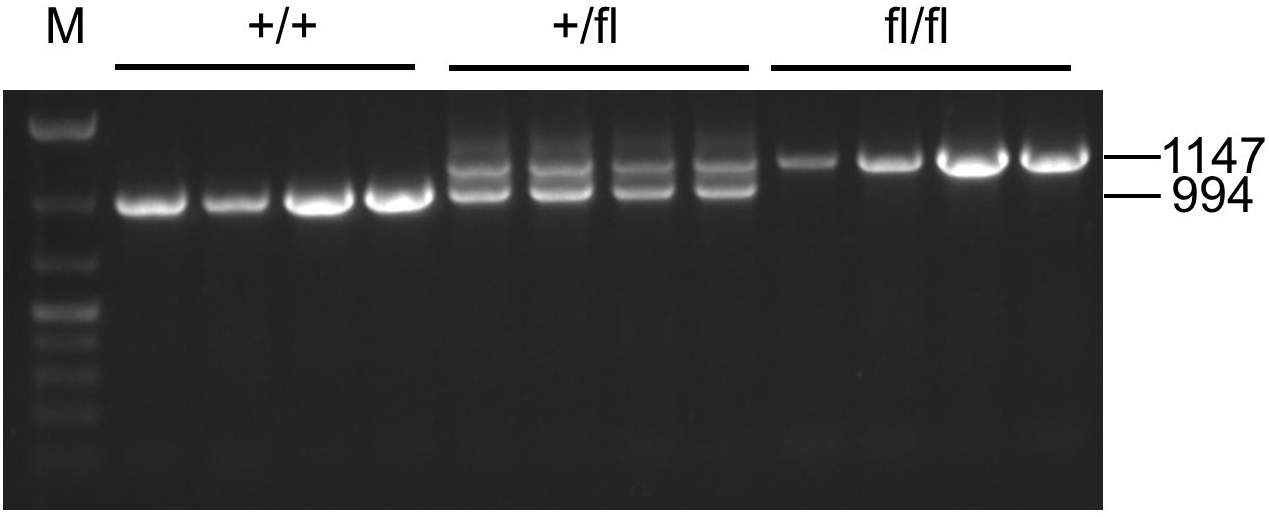
Absence of tamoxifen-inducible Cre-recombinase-based excision of the *Nos1ap* floxed allele in tail tissue of αMHC-MerCreMer mice. Post tamoxifen intraperitoneal injections, PCR amplification of the *Nos1ap* locus using tail tissue genomic DNA from *Nos1ap*^+/+^, *Nos1ap*^+/fl^ and *Nos1ap*^fl/fl^ mice, all with tamoxifen-inducible αMHC-MerCreMer transgene, shows absence of excision. M: DNA ladder.

**Figure S4:**
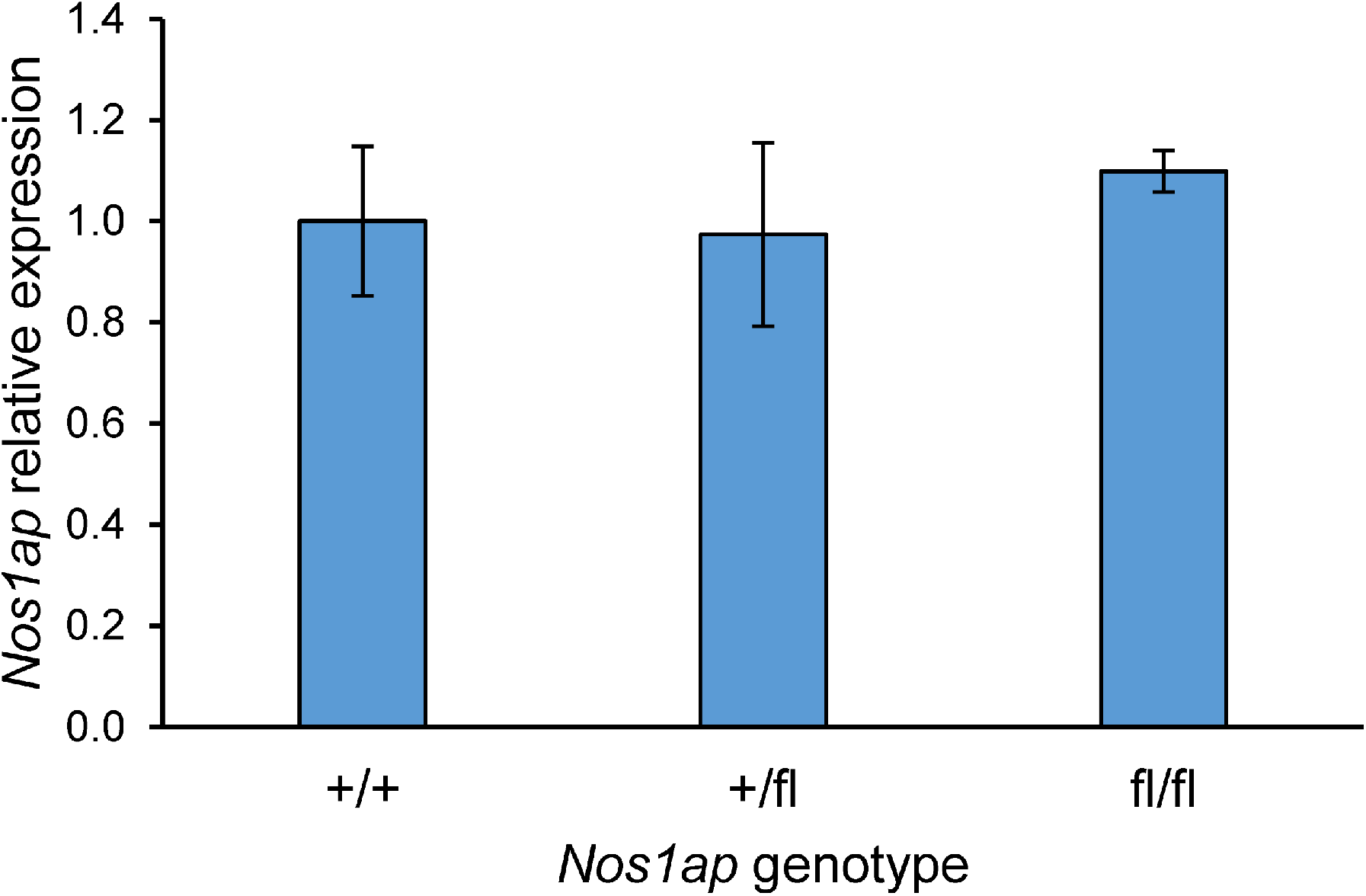
No significant differences in *Nos1ap* left ventricle gene expression in αMHC-MerCreMer-negative *Nos1ap* floxed allele carriers and homozygotes. In the absence of αMHC-MerCreMer, compared to wildtype mice (+/+), *Nos1ap* floxed (fl) allele carriers and homozygotes have no significant difference in *Nos1ap* expression in left ventricle tissue. Error bars: SEM; *n*=6 for each genotype.

**Figure S5:**
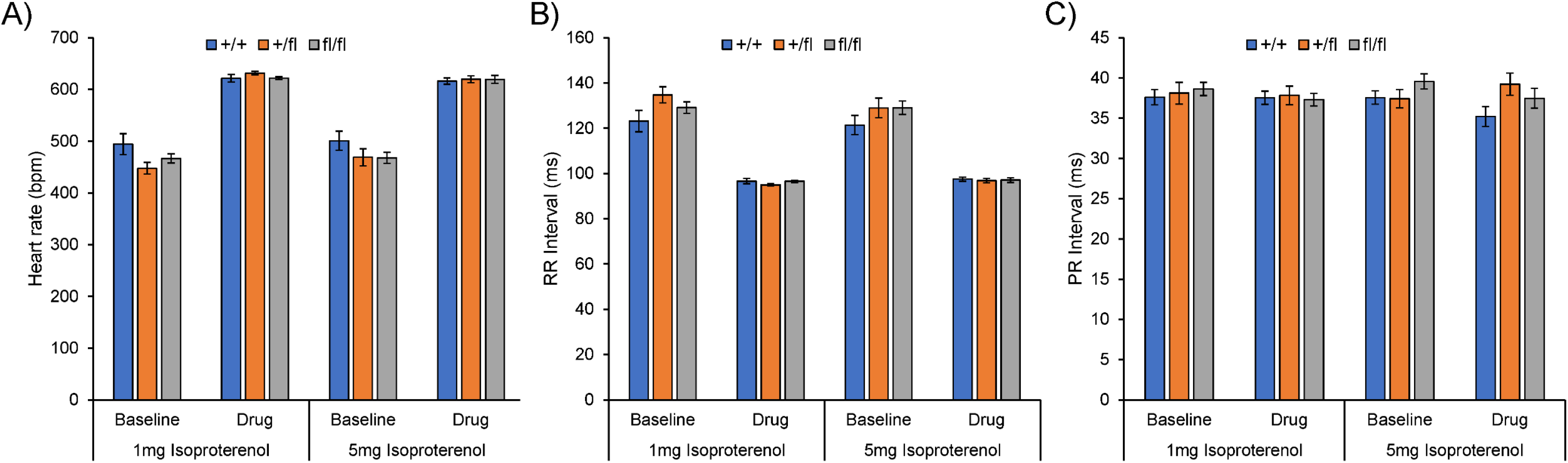
No significant differences in heart rate, RR interval and PR interval in anesthetized ECG measurements in *Nos1ap*^+/+^, *Nos1ap*^+/fl^ and *Nos1ap*^fl/fl^ mice with αMHC-MerCreMer. Heart rate (bpm: beats per minute) (*A*), RR interval (*B*) and PR interval (*C*) observed in surface ECG recordings of anesthetized mice at baseline and after injection of isoproterenol (1mg/kg or 5mg/kg dose). Error bars: SEM.

**Figure S6:**
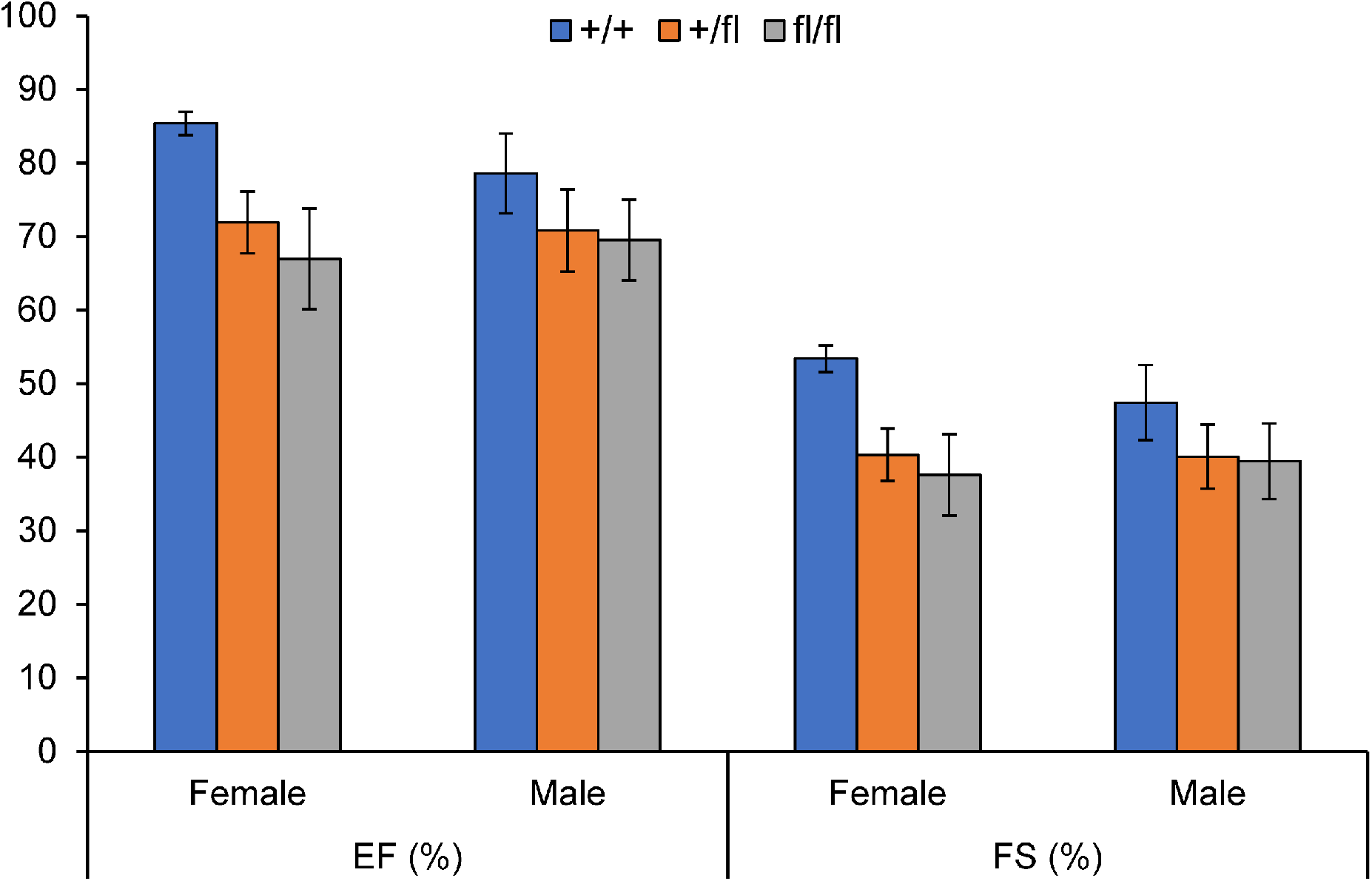
*Nos1ap* floxed allele carrier females with αMHC-MerCreMer trends towards reduced left ventricular function. Compared to wildtype mice, (+/+), *Nos1ap* floxed (fl) allele carrier and homozygote females show reduced ejection fraction (EF) and fractional shortening (FS) by echocardiography.

**Table S1:**
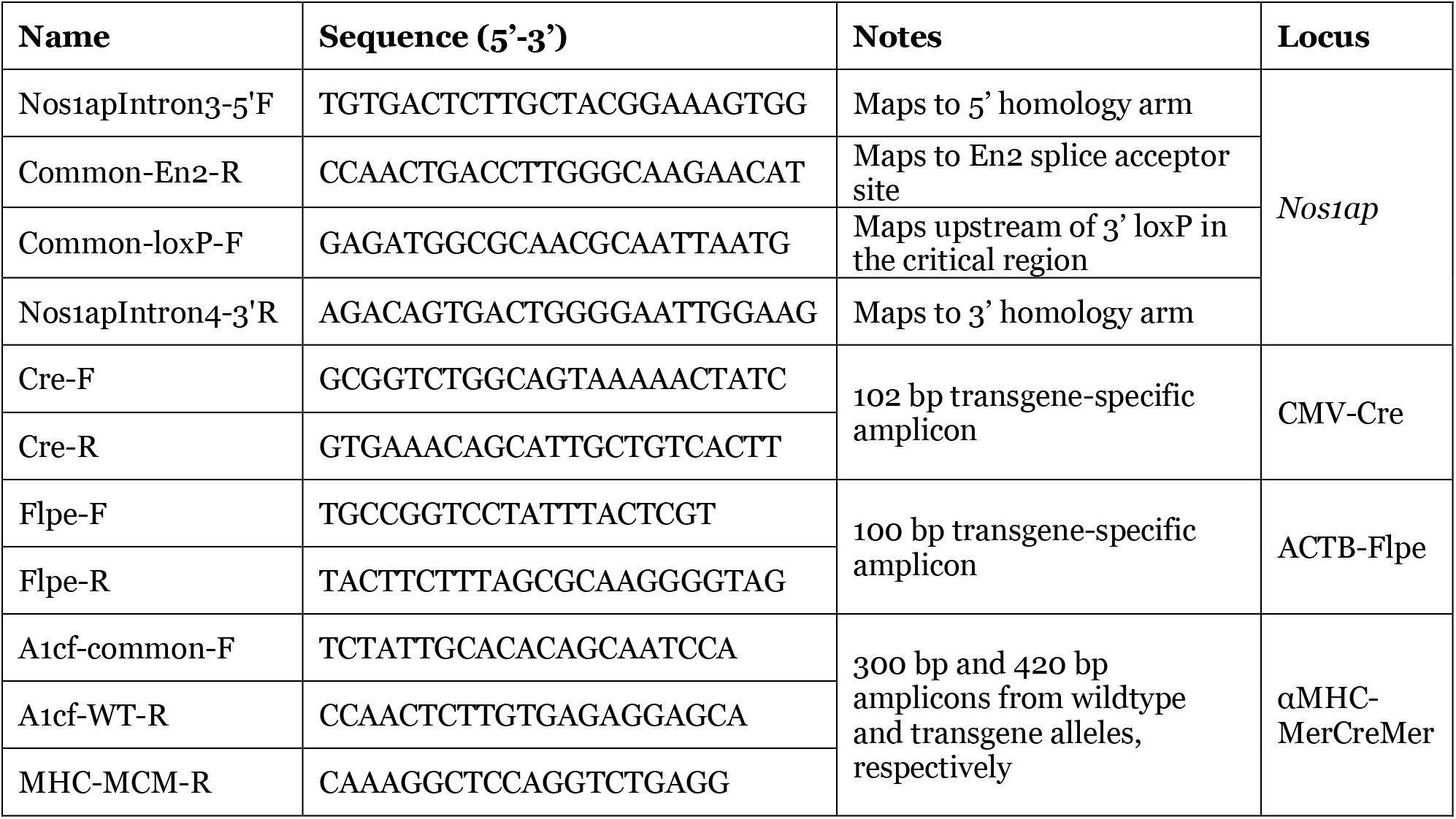
Primers for PCR genotyping.

**Table S2:**
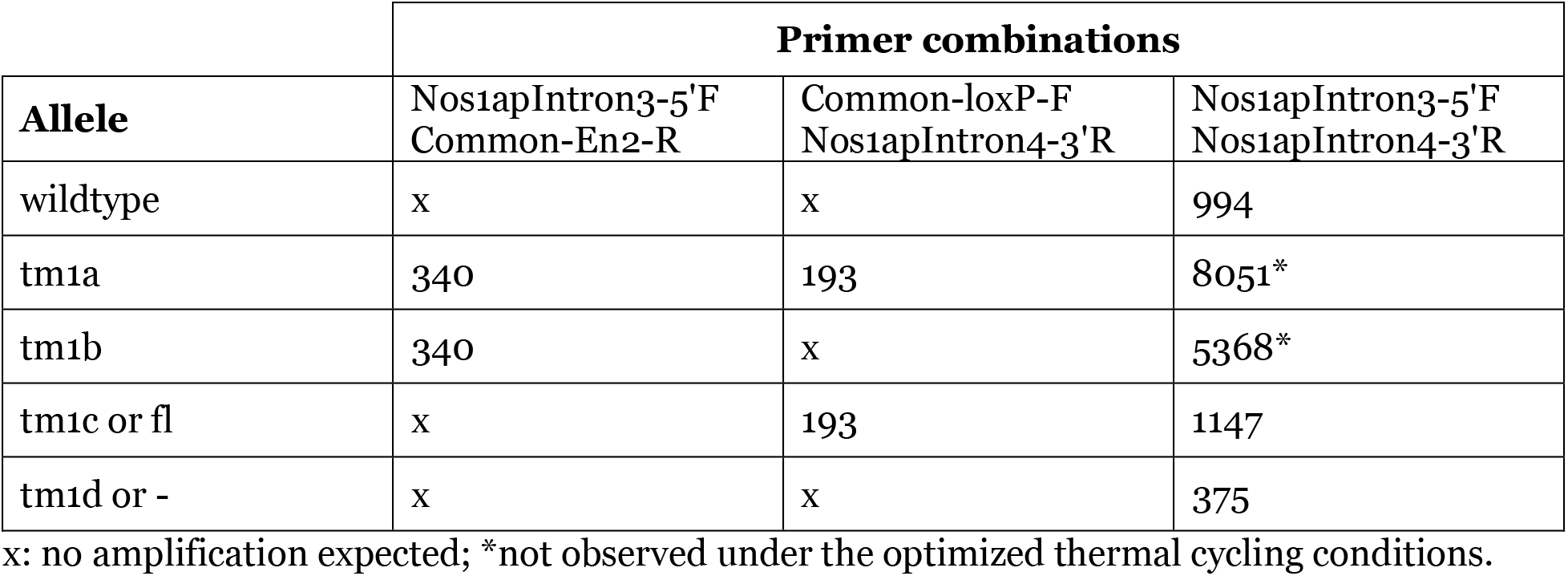
Alleles and PCR amplicon sizes (in bp)

**Table S3:** Genotype distributions from crosses between *Nos1ap*^+/fl^; +/+ and *Nos1ap*^+/fl^; +/*Tg*^αMHC-MerCreMer^ mice.

| Genotype | Observed (%) | Expected<sup>1</sup> (%) | $\chi^2$ | *P* |
| --- | --- | --- | --- | --- |
| *Nos1ap*<sup>+/+</sup>; +/+ | 14 (11.4) | 15 (12.5) | 2.69 | 0.75 |
| *Nos1ap*<sup>+/<sup>fl</sup></sup>; +/+ | 26 (21.1) | 31 (25.0) |  |  |
| *Nos1ap*<sup><sup>fl</sup>/<sup>fl</sup></sup>; +/+ | 17 (13.8) | 31 (12.5) |  |  |
| *Nos1ap*<sup>+/+</sup>; +/*Tg*<sup>αMHC-MerCreMer</sup> | 17 (13.8) | 15 (12.5) |  |  |
| *Nos1ap*<sup>+/<sup>fl</sup></sup>; +/*Tg*<sup>αMHC-MerCreMer</sup> | 29 (23.6) | 31 (25.0) |  |  |
| *Nos1ap*<sup><sup>fl</sup>/<sup>fl</sup></sup>; +/*Tg*<sup>αMHC-MerCreMer</sup> | 20 (16.3) | 15 (12.5) |  |  |

<sup>1</sup>rounded-off to single significant digit.

**Table S4:** Counts of animals per genotype and time-point undergoing awake ECG measurement.

|  | Time-point (weeks) |  |  |  |  |  |  |  |  |  |  |  |  |  |  |  |
| --- | --- | --- | --- | --- | --- | --- | --- | --- | --- | --- | --- | --- | --- | --- | --- | --- |
| Genotype | 6 | 8 | 10 | 12 | 14 | 16 | 18 | 20 | 22 | 24 | 28 | 32 | 36 | 48 | 44 | 48 |
| *Nos1ap*<sup>+/+</sup>;<br>+/αMHC-MCM | 15 | 15 | 14 | 14 | 12 | 11 | 11 | 11 | 11 | 11 | 11 | 11 | 11 | 11 | 11 | 11 |
| *Nos1ap*<sup>+/fl</sup>;<br>+/αMHC-MCM | 23 | 23 | 22 | 21 | 19 | 17 | 16 | 15 | 14 | 14 | 14 | 10 | 10 | 10 | 10 | 10 |
| *Nos1ap*<sup>fl/fl</sup>;<br>+/αMHC-MCM | 16 | 16 | 14 | 14 | 13 | 12 | 12 | 12 | 12 | 12 | 11 | 12 | 12 | 12 | 12 | 12 |
| *Nos1ap*<sup>+/+</sup>;<br>+/+ | 13 | 13 | 13 | 10 | 10 | 10 | 10 | 10 | 10 | 10 |  |  |  |  |  |  |
| *Nos1ap*<sup>+/fl</sup>;<br>+/+ | 26 | 26 | 24 | 23 | 23 | 21 | 18 | 17 | 16 | 13 |  |  |  |  |  |  |
| *Nos1ap*<sup>fl/fl</sup>;<br>+/+ | 13 | 14 | 12 | 12 | 11 | 10 | 10 | 10 | 10 | 9 |  |  |  |  |  |  |

**Table S5:** Age- and genotype-dependent effects on heart rate corrected QT interval observed in awake ECG measurement in *Nos1ap*^+/+^, *Nos1ap*^+/fl^ and *Nos1ap*^fl/fl^ mice with αMHC-MerCreMer.

| **Regression Statistics** |  |  |  |  |  |  |
| --- | --- | --- | --- | --- | --- | --- |
| Multiple R | 0.21 |  |  |  |  |  |
| R Square | 0.04 |  |  |  |  |  |
| Adjusted R Square | 0.04 |  |  |  |  |  |
| Standard Error | 1.91 |  |  |  |  |  |
| Observations | 1989 |  |  |  |  |  |
| **ANOVA** | df | SS | MS | F | Significance F |  |
| Regression | 3 | 339.74 | 113.25 | 31.15 | 1.13E-19 |  |
| Residual | 1985 | 7217.31 | 3.64 |  |  |  |
| Total | 1988 | 7557.04 |  |  |  |  |
| **Predictors** | Coefficients | Standard Error | t Stat | P-value | Lower 95% | Upper 95% |
| Intercept | 45.42 | 0.11 | 415.59 | 0 | 45.21 | 45.64 |
| Age weeks | 0.03 | 0.00 | 9.08 | 2.48E-19 | 0.02 | 0.04 |
| (fl+) | 0.10 | 0.10 | 0.91 | 0.363 | -0.11 | 0.30 |
| (fl/fl) | 0.31 | 0.11 | 2.84 | 0.005 | 0.10 | 0.52 |

**Table S6:** Age- and genotype-dependent effects on heart rate corrected QT interval observed in awake ECG measurement in *Nos1ap*^+/+^, *Nos1ap*^+/fl^ and *Nos1ap*^fl/fl^ mice without αMHC-MerCreMer.

|  |  |  |  |  |  |  |
| --- | --- | --- | --- | --- | --- | --- |
| **Regression Statistics** |  |  |  |  |  |  |
| Multiple R | 0.18 |  |  |  |  |  |
| R Square | 0.03 |  |  |  |  |  |
| Adjusted R Square | 0.03 |  |  |  |  |  |
| Standard Error | 1.97 |  |  |  |  |  |
| Observations | 1345 |  |  |  |  |  |
| **ANOVA** | df | SS | MS | F | Significance F |  |
| Regression | 3 | 170.27 | 56.76 | 14.64 | 2.20E-09 |  |
| Residual | 1341 | 5199.21 | 3.88 |  |  |  |
| Total | 1344 | 5369.48 |  |  |  |  |
| **Predictors** | Coefficients | Standard Error | t Stat | P-value | Lower 95% | Upper 95% |
| Intercept | 44.93 | 0.17 | 264.13 | 0 | 44.59 | 45.26 |
| Age weeks | 0.04 | 0.01 | 4.27 | 2.11E-05 | 0.02 | 0.06 |
| (fl/+) | 0.65 | 0.13 | 4.97 | 7.65E-07 | 0.39 | 0.91 |
| (fl/fl) | 0.60 | 0.15 | 4.02 | 6.16E-05 | 0.31 | 0.90 |

**Table S7:** Genotype-, isoproterenol dose- and condition-dependent effects on QRS interval in anesthetized ECG measurement in *Nos1ap*^+/+^ and *Nos1ap*^fl/fl^ mice with αMHC-MerCreMer.

|  |  |  |  |  |  |  |
| --- | --- | --- | --- | --- | --- | --- |
| **Regression Statistics** |  |  |  |  |  |  |
| Multiple R | 0.69 |  |  |  |  |  |
| R Square | 0.47 |  |  |  |  |  |
| Adjusted R Square | 0.45 |  |  |  |  |  |
| Standard Error | 0.89 |  |  |  |  |  |
| Observations | 86 |  |  |  |  |  |
| **ANOVA** | df | SS | MS | F | Significance F |  |
| Regression | 3 | 58.18 | 19.39 | 24.24 | 2.51E-11 |  |
| Residual | 82 | 65.61 | 0.80 |  |  |  |
| Total | 85 | 123.79 |  |  |  |  |
| **Predictors** | Coefficients | Standard Error | t Stat | P-value | Lower 95% | Upper 95% |
| Intercept | 8.61 | 0.23 | 38.23 | 4.91E-54 | 8.16 | 9.06 |
| condition | 1.44 | 0.19 | 7.47 | 7.75E-11 | 1.06 | 1.82 |
| dose | -0.04 | 0.05 | -0.74 | 0.46 | -0.13 | 0.06 |
| genotype | -0.78 | 0.19 | -4.03 | 1.25E-04 | -1.16 | -0.39 |

